# Single-fluorescent protein reporters allow parallel quantification of NK cell-mediated granzyme and caspase activities in single target cells

**DOI:** 10.1101/328260

**Authors:** Clarissa Liesche, Patricia Sauer, Maren Claus, Roland Eils, Joël Beaudouin, Carsten Watzl

## Abstract

Natural killer (NK) cells eliminate infected and tumorigenic cells through delivery of granzymes via perforin pores or by activation of caspases via death receptors. In order to understand how NK cells combine different cell death mechanisms it is important to quantify target cell responses on a single cell level. However, currently existing reporters do not allow the measurement of several protease activities inside the same cell. Here we present a strategy for the comparison of two different proteases at a time inside individual target cells upon engagement by NK cells. We developed single-fluorescent protein reporters containing the RIEAD or the VGPD cleavage site for the measurement of granzyme B activity. We show that these two granzyme B reporters can be applied in combination with caspase-8 or caspase-3 reporters. While we did not find that caspase-8 was activated by granzyme B, our method revealed that caspase-3 activity follows granzyme B activity with a delay of about 6 minutes. Finally, we illustrate the comparison of several different reporters for granzyme A, M, K and H. The here presented approach is a valuable means for the investigation of the temporal evolution of cell death mediated by cytotoxic lymphocytes.

## 2. Introduction

As part of the innate immune system, Natural Killer (NK) cells can eliminate infected and tumorigenic cells (Watzl, 2014). To do so, they adhere to a target cell and establish an immunological synapse (Mace et al., 2014). The following NK cell receptor signaling can trigger the release of cytotoxic granules from the NK cell into the cleft of this synapse (Watzl et al., 2014). Like cytotoxic T lymphocytes (CTLs), NK cells have two mechanisms to induce cell death of target cells (Ewen et al., 2012; Strasser et al., 2009): In the first mechanism, granzymes are released from cytotoxic granules and enter the target cell via perforin pores and induce cell death. In the second mechanism, CD95L or TRAIL are presented at the surface of NK cells and induce extrinsic apoptosis in target cells through activation of the death receptors CD95 or TRAIL-R1/-R2.

How NK cells orchestrate the activities of granzymes and the activation of extrinsic apoptosis remains poorly understood. Extrinsic apoptosis starts with the formation of the so-called death inducing signaling complex (DISC), composed of activated death receptors and recruited FADD adaptor proteins and initiator procaspases-8/-10. Once activated, these caspases cleave and activate effector procaspase-3/-7 (Peter and Krammer, 2003; Stennicke et al., 1998), leading to apoptosis, unless presence of XIAP blocks their activity (Barnhart et al., 2003; Wowk and Trapani, 2004). When the pro-apoptotic Bcl-2 protein BID is cleaved by caspase-8/-10 in sufficient amount, truncated BID induces mitochondrial outer membrane permeabilization. Subsequent release of cytochrome c activates caspase-9, while release of SMAC induces the degradation of XIAP, both leading to massive activation of effector caspases.

To deliver granzymes in the cytosol of target cells, perforin forms a pore in cellular membranes (Law et al., 2010). It is debated if this occurs at the plasma membrane (Lopez et al., 2013a, 2013b) or the membrane of endosomes (Browne et al., 1999; Froelich et al., 1996; Thiery et al., 2010). Of the five human granzymes A, B, H, K and M, granzyme B is the best characterized one and shares substrate specificity with caspases for cleavage after aspartate residues (Bots and Medema, 2006; Joeckel and Bird, 2014; Susanto et al., 2012). Both, granzyme B and caspase-8 can cleave BID, yet at different sites, at D75 (RIEAD’S) and D60 (ELQTD’G) (Li et al., 1998), respectively. While granzyme B has been shown to cleave the initiator procaspase-8 (Medema et al., 1997) and the effector procaspase-3 (Quan et al., 1996)(Yang *et al*, 1998; Andrade *et al*, 1998; Atkinson *et al*, 1998a)(Goping et al., 2003), other substrates measured *in vitro* have been reported to be more efficiently cleaved, for example DNA-PKc or BID (Andrade et al., 1998; Barry et al., 2000b; Pinkoski et al., 2001; Sutton et al., 2000). From this perspective, granzyme B is suggested to play a role not only as an initiator but also as executioner enzyme in target cell death (Wowk and Trapani, 2004).

Having reporters that would allow the measurement of the contribution of granzymes and caspases in a single cell would be beneficial to characterize the activity of NK cells. Specific protease biosensors based on luciferase (Li et al., 2014; Vrazo et al., 2015), fluorophore quenching (Packard et al., 2007) and FRET (Choi and Mitchison, 2013; Zhu et al., 2016) (**Table 1**) have facilitated the study of the killing mechanism by granzymes and death receptors. However they do not easily allow multiplexing for the quantification of several protease activities in single cells. Parallel assessment of protease activity inside single cells would allow for a better understanding of the temporal order of signaling events in the NK cell killing mechanism. In order to reach this aim, we present an approach to measure NK cell-mediated activity of two proteases at once in single target cells. We demonstrate our approach by measuring granzyme B, caspase-8 and caspase-3 activity in target cells exposed to NK cells. The pallet of reporters can easily be extended by cloning cleavage linkers, as illustrated here with the measurement of potential substrates for different granzymes. We believe that these reporters offer a valuable resource to characterize the physiology of NK cells or to test the activity of patient-derived NK cells.

**Table 1:**
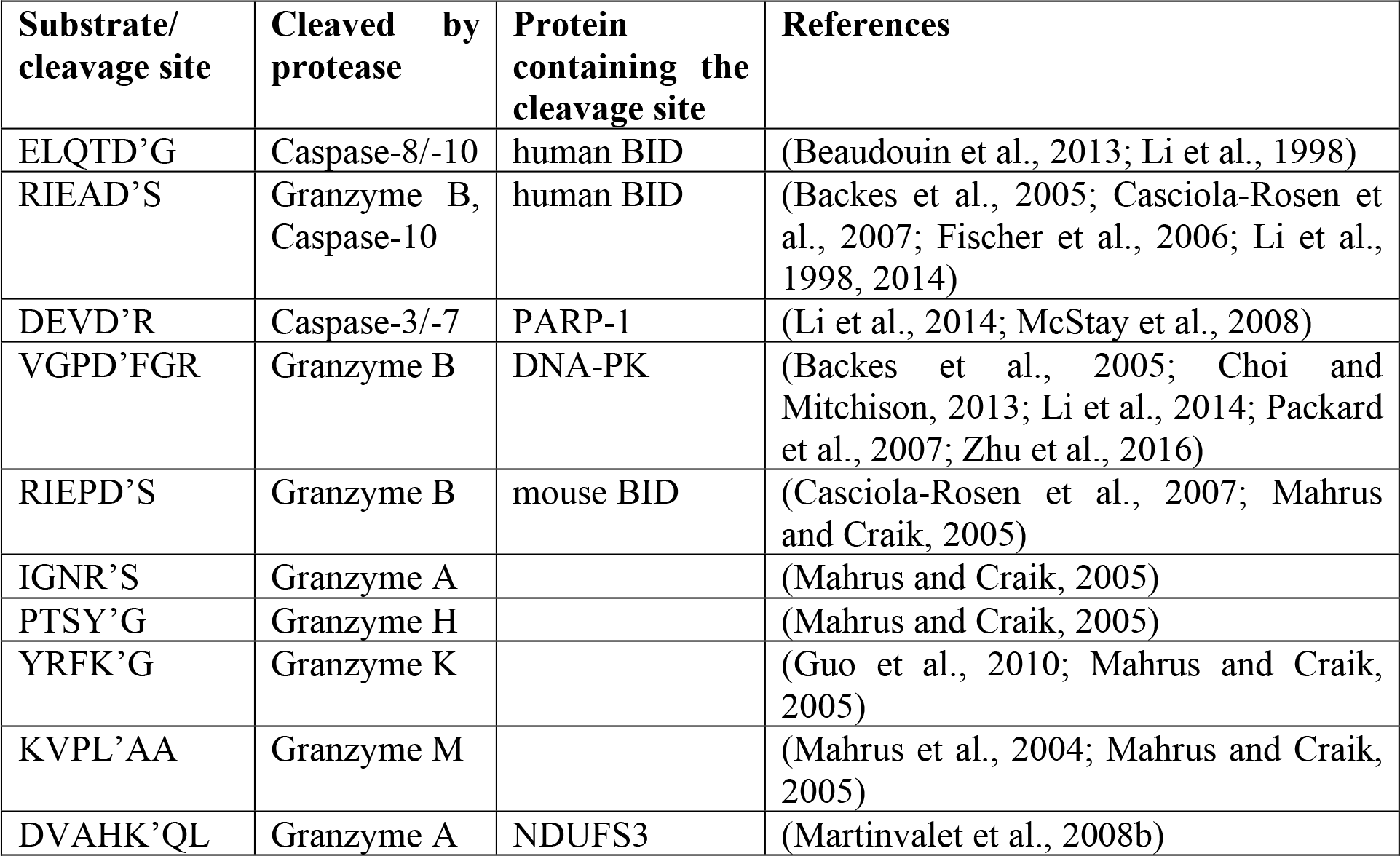
Cleavage sites used in this study.

### 3. Results

#### 3.1. Measurement of protease activity induced by NK cells in living target cell

To get insights into the process by which individual NK cells kill their target cells, we aimed at comparing the strength of different proteases inside single target cells. To achieve this, we designed fluorescent reporters following the strategy that we previously established to measure caspase activity upon CD95 activation (Beaudouin et al., 2013). These reporters consist of one fluorescent protein fused to a localization domain through a linker that can be specifically cleaved by proteases. In this study we use the nuclear export signal (NES) as localization domain (**Fig. 1**). As exemplified below, this reporter allows an easy quantification of protease activity, and it detects any activity originating from an enzyme facing the cytosol. Protease cleavage leads to separation of the fluorescent protein from the localization domain. The free fluorescent protein is small enough to enter or exit the nucleus by passive diffusion, a process that takes about 1 min. Reporter cleavage can be quantified by measuring the increase of fluorescence signal in the nucleus. This is ideally imaged by time-lapse fluorescence confocal microscopy as this allows the measurement of the fluorescence intensity inside the nucleus without contamination of signal from the cytosol below and above. Thus, by quantifying the spatial redistribution of the fluorescence signal inside the target cell, reporter cleavage can be calculated. Image analysis can be done using the freely available ImageJ software and proceeds as follows: generation of an image stack from time series data, background subtraction and measurement of the mean fluorescence signal in a region of interest representing the cell nucleus. To estimate the extent of substrate cleavage, this nuclear signal is normalized to the cytosolic one. One possibility is to perform this normalization for each time point, which requires more work but has the advantage of correcting for potential photobleaching over time:

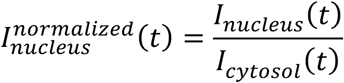

**Fig. 1:**
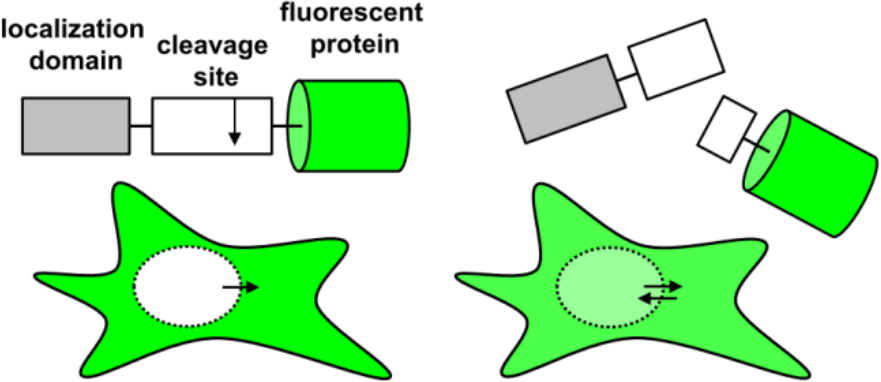
Schematic representation of a single-fluorescent protein (single FP) reporter and its fluorescence distribution inside the cell when using a nuclear export signal as localization domain. Upon cleavage of the reporter, the fluorescent protein can diffuse passively into the nucleus.

In this case, the normalized intensity tends towards 1, or 100%, when cleavage is complete. The here presented data were analyzed in this way. The second possibility consists of normalizing with the cytosolic signal before the addition of the NK cells. This approach is relevant if photobleaching can be neglected:

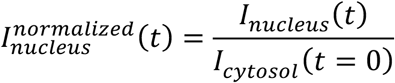

In this case, as the nucleus and the cytoplasm are roughly occupying the same volume, the normalized intensity tends towards 0.5, or 50%.

Since each reporter contains only one fluorescent protein, these so-called single-fluorescent protein reporters allow parallel assessment of several reporters within the same cell. Hence, different comparison can be realized: the measurement of (i) different cleavage sites for different proteases or of (ii) different cleavage sites for the same protease.

#### 3.2. Granzyme B activity can be measured with single-fluorescent protein reporters having the RIEADS or the VGPD cleavage site

In order to establish single-fluorescent protein reporters for the measurement of granzyme B activity, we designed and tested two different reporters carrying a linker sequence known to be cleaved by granzyme B. The first reporter carries the amino acid sequence RIEADS (single amino acid code), which is present in the protein BID. The second reporter carries the amino acid sequence VGPD from the protein DNA-PKc as cleavable linker. We co-expressed the two reporters NES-RIEADS-mCherry and NES-VGPD-mGFP in HeLa cells expressing CD48, a ligand for the activating NK cell receptor 2B4 (CD244), which renders them more sensitive to killing by NK cells (Hoffmann et al., 2011). The transfected target cells were imaged by confocal microscopy over time upon addition of the human NK cell line NK92-C1. Cell death was recognized from images by cell rounding and cell shrinkage following NK cell engagement (**Fig 2A**). About 20 minutes before target cell death, NES-RIEADS-mCherry and NES-VGPD-mGFP reporter cleavage was detected from the appearance of fluorescence inside the nucleus (**Fig 2A**). In order to plot the cleavage kinetics of several cells independently of the time of NK cell engagement, we defined the time of death as time point zero. In this way, we found that both reporters were cleaved on average at the same rate before target cell death (**Fig 2B**). Furthermore, the cleavage efficiency of NES-RIEADS-mCherry and NES-VGPD-mGFP reporters inside the same cell correlated well, showing that both reporters are able to mirror the observed cell-to-cell variability of granzyme B activity (**Fig 2B**). We next verified the specificity of these two reporters by measuring their cleavage in cells that overexpress serpin B9, a natural inhibitor of granzyme B (Kaiserman and Bird, 2010). Compared to control cells, the time of cell death was delayed in serpin B9 overexpressing cells (Fig 2C **and D**). Consistent with that, reporter cleavage was reduced from 53% to 17% (NES-RIEADS-mCherry) and from 34% to 6% (NES-VGPD-mCherry), showing that both reporters are suitable for the specific measurement of granzyme B activity.

**Fig. 2:**
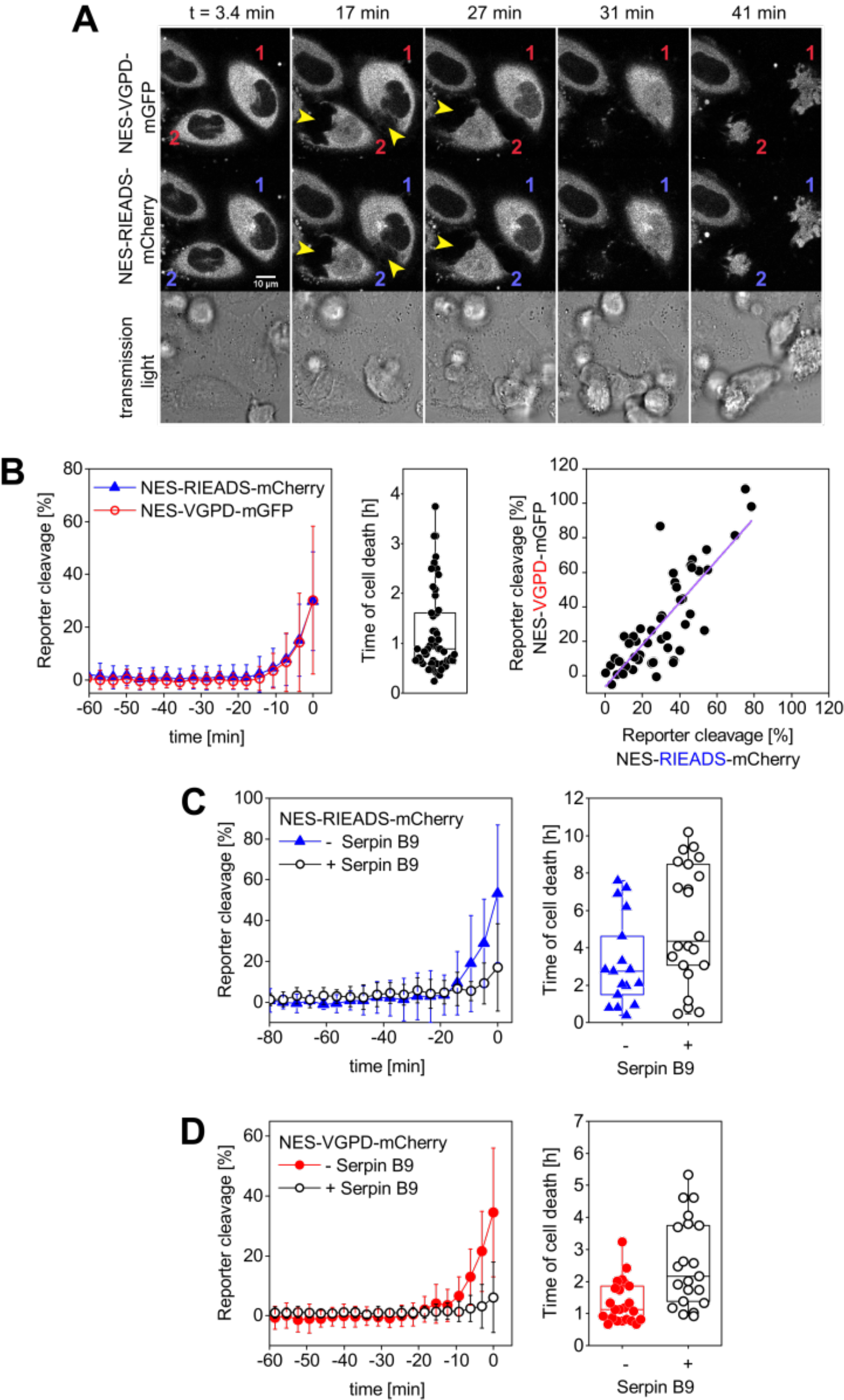
Measurement of granzyme B activity in target cells using single FP-reporters. (**A-B**) HeLa cells transfected with CD48, NES-VGPD-mGFP and NES-RIEADS-mCherry were imaged by confocal microscopy upon incubation with 3-fold more NK92-C1 cells than target cells. (**A**) Representative images from confocal microscopy. Two target cells were labelled 1 and 2. Their contact with NK cells can be seen on fluorescence images (arrowheads). Appearance of fluorescence signal in the nucleus shows reporter cleavage. (**B**) NES-VGPD-mGFP and NES-RIEADS-mCherry reporter cleavage quantification. The last measurement point preceding cell death was set to time = 0 in order to calculate the mean and standard deviation from 51 single cell measurements as shown. The boxplot shows the time of cell death of each cell. The scatter plot shows the correlation between NES-VGPD-mGFP and NES-RIEADS-mCherry reporter cleavage. Values correspond to the measurement point preceding cell death. Linear fit of data with slope =1.23 ± 0.12 and Pearson’s correlation coefficient = 0.82. (**C-D**) SerpinB9 expression in target cells reduces NES-VGPD-mCherry and NES-RIEADS-mCherry reporter cleavage induced by NK92-C1 cells. (**C**) HeLa(CD48) cells were transfected with NES-RIEADS-mCherry reporter alone or with the NES-RIEADS-mCherry reporter and serpin B9, E:T ratio = 1. Mean and standard deviation of 17 and 22 cells, respectively. (**D**) HeLa(CD48) cells were transfected with NES-VGPD-mCherry reporter alone or with the NES-VGPD-mCherry reporter and serpin B9, E:T ratio = 3. Mean and standard deviation of 23 and 22 cells, respectively. For cell death times shown in boxplots C and D, the mean between the control group and the test group (cells transfected with serpinB9) is significantly different at the 0.05 level.

#### 3.3. Single-fluorescent protein reporters allow multiplexing: distinguishing different proteases inside the same cell

CD95 signaling in HeLa cells leads to notable caspase-8 and -3 activation (Beaudouin et al., 2013). On the one hand it was reported that caspase-8 can get activated through cleavage by granzyme B (Medema et al., 1997). On the other hand it was shown that granzyme B activates effector caspases through cleavage of BID (Atkinson et al., 1998b; Goping et al., 2003; Pinkoski et al., 2001; Sutton et al., 2000). Deciphering the contribution of caspase-8 and granzyme B to NK cell mediated cell death, and in particular the potential activation of caspase-8 by granzyme B, is an intriguing question that could help to better understand cell death signaling by NK cells.

Therefore, we tested if caspase-8 and granzyme B activity can be distinguished within the same cell using our single-fluorescent protein reporters. For this aim, we measured NES-ELQTD-mGFP (for caspase-8), together with either NES-RIEADS-mCherry or NES-VGPD-mCherry (for granzyme B). Upon addition of soluble trimerized CD95L (IZsCD95L) to HeLa cells, we observed efficient cleavage of the caspase-8 reporter NES-ELQTD-mGFP. In contrast, in the same cells, the NES-RIEADS-mCherry reporter was cleaved only up to 10% (**Fig 3A**) and the NES-VGPD-mCherry reporter was virtually not cleaved (**Fig 3B**). This shows that granzyme B activity (using either the RIEADS-or the VGPD-reporter) can be clearly distinguished from caspase-8 activity (using the ELQTD-reporter) within the same cell.

**Fig. 3:**
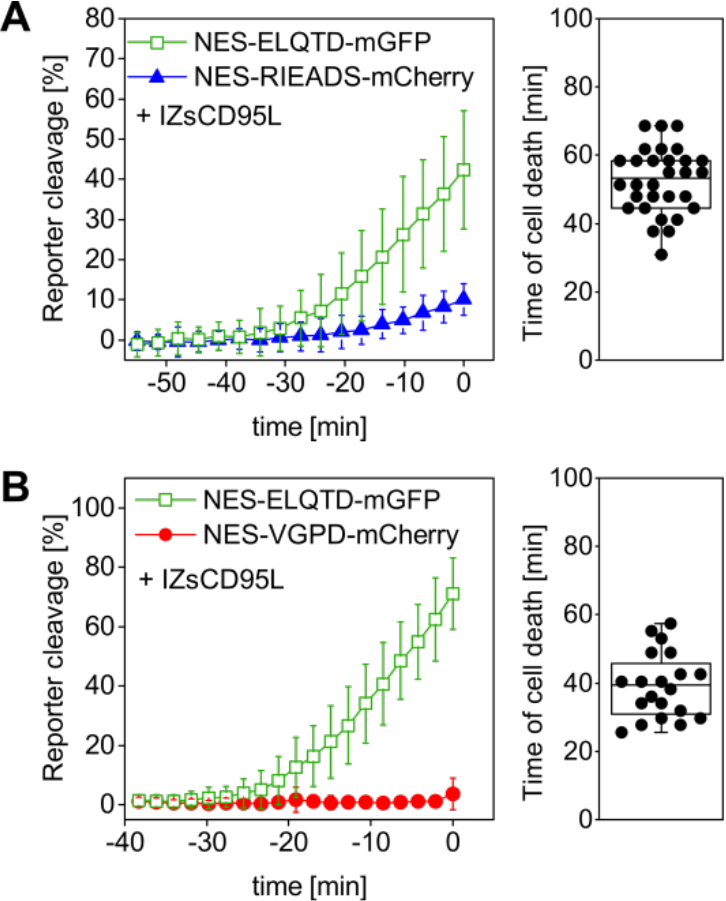
Granzyme B activity can be distinguished from caspase-8 activity. HeLa cells overexpressing CD95 were transfected with (**A**) NES-ELQTD-mGFP (for caspase-8) and NES-RIEADS-mCherry for granzyme B or (**B**) NES-ELQTD-mGFP for caspase-8 and NES-VGPD-mCherry for granzyme B and imaged by confocal microscopy upon incubation with 1 μg/ml IZsCD95L.

We next set out to test for caspase-8 activity in cells showing granzyme B activity. For this, we compared reporter cleavage in HeLa cells in the absence or presence of CD95 expression. Transient expression of shRNA against CD95 led to absence of cell death and caspase-8 activity in a control experiment using IZsCD95L as inducer (**Fig. 4A**). The nuclear protein H2B-eBFP2 served as a fluorescent marker to recognize cells that contain the plasmid encoding shRNA (**Fig. 4B**). Upon addition of NK cells to HeLa(CD48) cells expressing control shRNA, we observed NES-RIEADS-mCherry reporter cleavage as expected indicating granzyme B activity. In addition, the NES-ELQTD-mGFP reporter was cleaved on average up to 7% indicating an additional activation of caspase-8 (**Fig 4C**). This caspase-8 activity could be due to the activation of caspase-8 by granzyme B, or due to the activation of death receptors during the engagement by the NK cells. To distinguish these two possibilities, we repeated the experiment in HeLa cells expressing a shRNA against CD95. The absence of CD95 expression abolished NES-ELQTD-mGFP reporter cleavage (**Fig 4D**). This demonstrates that the caspase-8 activity was due to activation of the CD95 pathway by NK cells and it also reveals that caspase-8 is not, or not efficiently, activated by granzyme B.

**Fig. 4:**
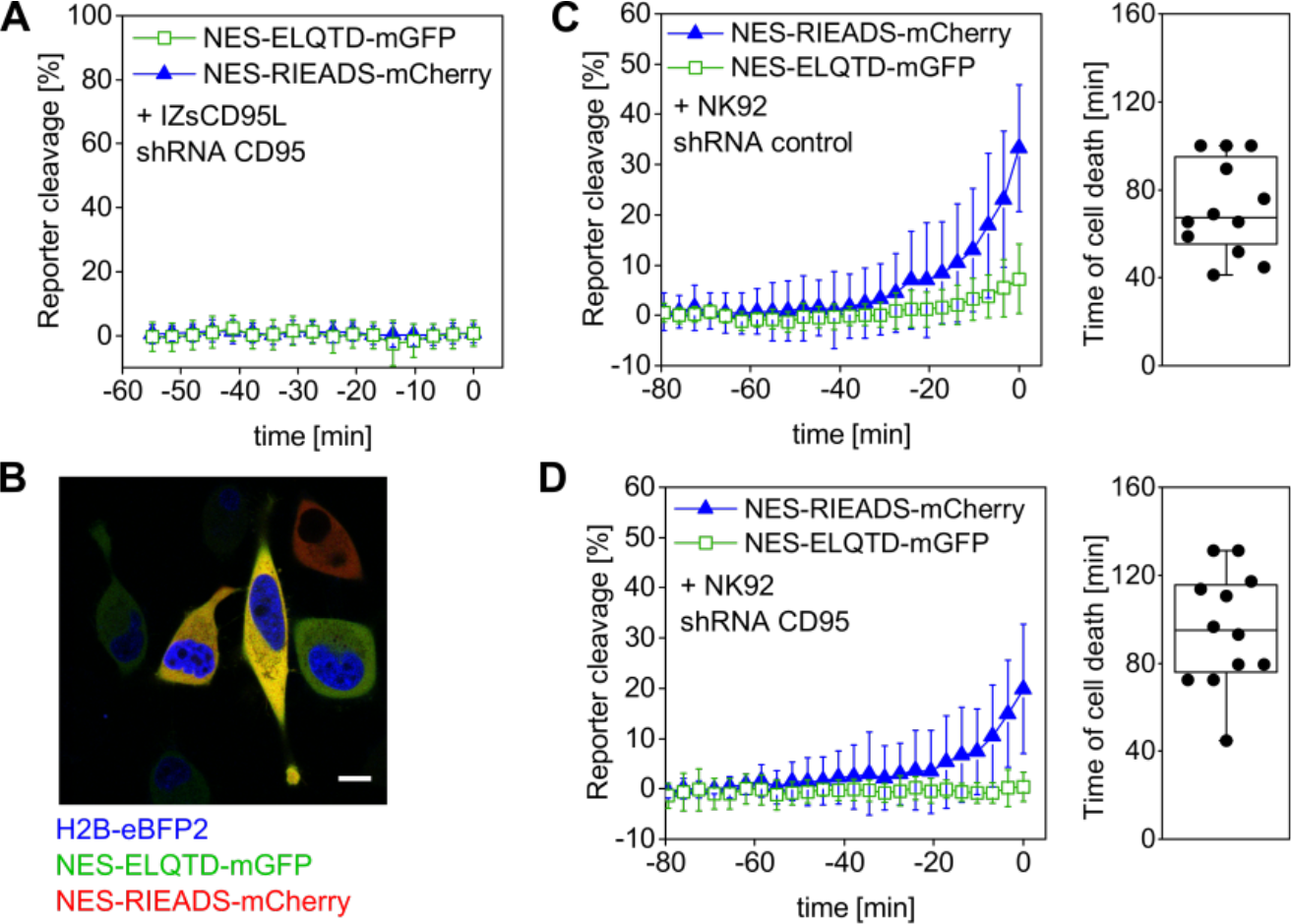
Caspase-8 activity generated by NK92 cells is not induced by granzyme B. HeLa(CD48) cells were triple-transfected with NES-ELQTD-mGFP (for caspase-8) and NES-RIEADS-mCherry (for granzyme B) together with a plasmid encoding shRNA against CD95 and the fluorescent marker H2B-eBFP2 (A, D) or with a plasmid encoding control shRNA and H2B-eBFP2 (C). (**A**) Cells expressing H2B-eBFP2 (and shRNA against CD95) and incubated with 1 μg/ml IZsCD95L neither showed reporter cleavage nor cell death. The plot shows data from 10 analyzed cells with time zero being the end of the microscopy time series. (**B**) Field of view showing HeLa(CD48) cells imaged by confocal fluorescence microscopy. The image is an overlay of three fluorescence emission channels: The nuclear protein H2B-eBFP2 (blue) was expressed as fluorescent marker from the same plasmid encoding shRNA against CD95. NES-ELQTD-mGFP (green) and NES-RIEADS-mCherry (red) overlay result in yellow color depending on reporter amounts. Scale bar: 10 μm. (**C**) Cells expressing control shRNA and co-cultured with NK92-C1 cells (E:T = 1) showed strong NES-RIEADS-mCherry reporter cleavage. The same cells showed on average up to 7% NES-ELQTD-mGFP reporter cleavage. (**D**) NES-ELQTD-mGFP reporter cleavage was absent in cells expressing shRNA against CD95, while granzyme B activity was clearly present in the same cells as seen from NES-RIEADS-mCherry reporter cleavage.

It is known that effector caspase-3 activation can be a consequence of granzyme B activity (Barry et al., 2000a). We thus hypothesized that caspase-3 activity should occur with a delay after granzyme B activity, as in the case of CD95 mediated extrinsic apoptosis (Beaudouin et al., 2013; Stennicke et al., 1998). To test this, we measured reporter cleavage of NES-VGPD-mCherry for granzyme B and NES-DEVDR-GFP for caspase-3 in HeLa cells upon addition of NK cells (**Fig 5A**). We found that caspase-3 activity appears on average about 6 minutes later than granzyme B (**Fig 5B**). The onset of caspase-3 activity was about 10 minutes before cell death, highlighting the speed of this cellular process. These data also show how precisely cell death can be analyzed on the single cell level with this approach using single-fluorescent protein reporters.

**Fig. 5:**
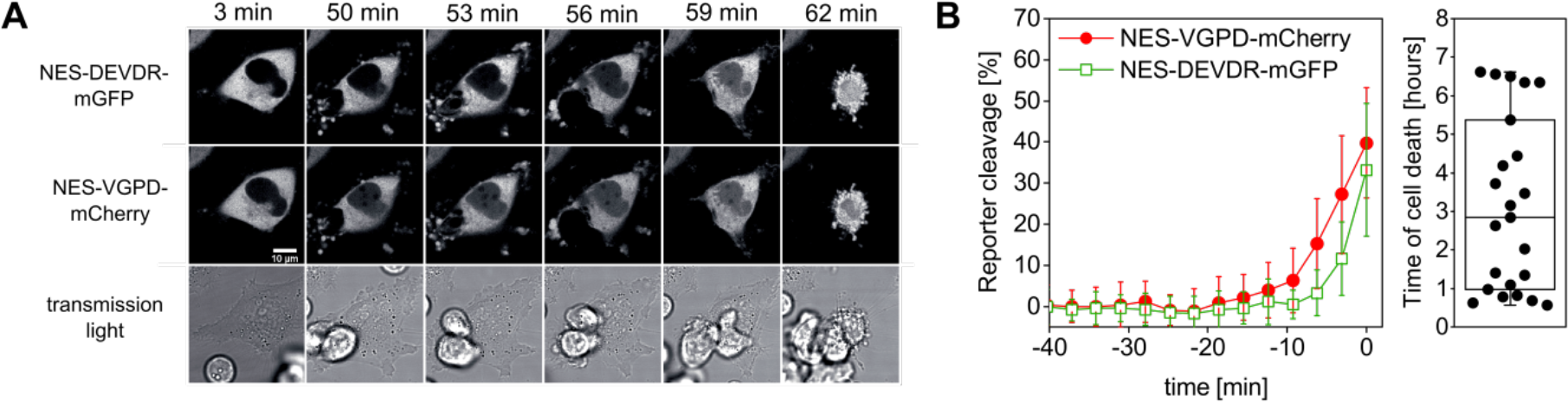
Measurement of granzyme B and caspase-3 activity in single target cells. (**A, B**) HeLa cells transfected with CD48, NES-VGPD-mCherry (for granzyme B) and NES-DEVDR-mGFP (for caspase-3) were imaged by confocal microscopy upon incubation with NK92-C1 cells (E:T = 2). (**A**) Example images of the time series. Note that the fluorescence signal inside the nucleus is visible for NES-VGPD-mCherry at t = 50 min and for NES-DEVDR-mGFP at t = 56 min. Scale bar: 10 μm. (**B**) Quantification of reporter cleavage of 23 cells and boxplot showing the time of target cell death. On average, caspase-3 activity appears with a short delay of about 6 min after granzyme B activity.

#### 3.4. Testing potential reporters for different granzymes

NK cells notably express granzyme A and B (Bratke et al., 2005; Grossman et al., 2004) but they also express the less characterized granzymes H, K and M (Krzewski and Coligan, 2012). In order to further illustrate our approach and to potentially capture the activity of granzymes A, M, H and K in single target cells, we designed fluorescent reporters that contain cleavage site candidates based on existing data from literature (Table 1). Using positional scanning combinatorial libraries optimal substrates for the five human granzymes have been previously identified and characterized (Mahrus and Craik, 2005). Among them, the IEPD and IGNR sequence have been further developed as fluorescent label and inhibitor for granzyme B and granzyme A (Mahrus and Craik, 2005). Moreover, optimal substrates were determined to be YRFK for granzyme K, KVPL for granzyme M and PTSY for granzyme H (Mahrus and Craik, 2005). In this study, we additionally designed a reporter containing the sequence DVAHKQL, which is present in the mitochondrial complex I protein NDUFS3, since granzyme A was reported to cleave this protein at this site after the amino acid lysine in this sequence (Martinvalet et al., 2008a). We measured these six different reporters in comparison to the granzyme B reporters NES-VGPD-mCherry or NES-RIEADS-mGFP. We found that the RIEADS and VGPD reporters were more sensitive to detect granzyme B activity compared to the IEDP reporter **(Fig. 6A).** Also, granzyme B activity was clearly dominant among the different experiments as at the time of death IGNRS (granzyme A), PTSYG (granzyme H), and YRFKG (granzyme K) reporter cleavage was in almost all cases less than RIEADS or VGPD (granzyme B) cleavage (**Fig. 6**). Moreover, no cleavage of DVAHQL (granzyme A) and KVPL (granzyme M) reporters could be observed. This comparison of different reporters would suggest that granzyme B and granzyme A are the most abundant granzymes used by the NK92-C1 to cleave substrates in the cytosol of target cells. This experiment shows how single-fluorescent protein reporter can be used to examine protease activity that is either induced inside target cells or stemming from cytotoxic lymphocytes such as NK cells.

**Fig. 6:**
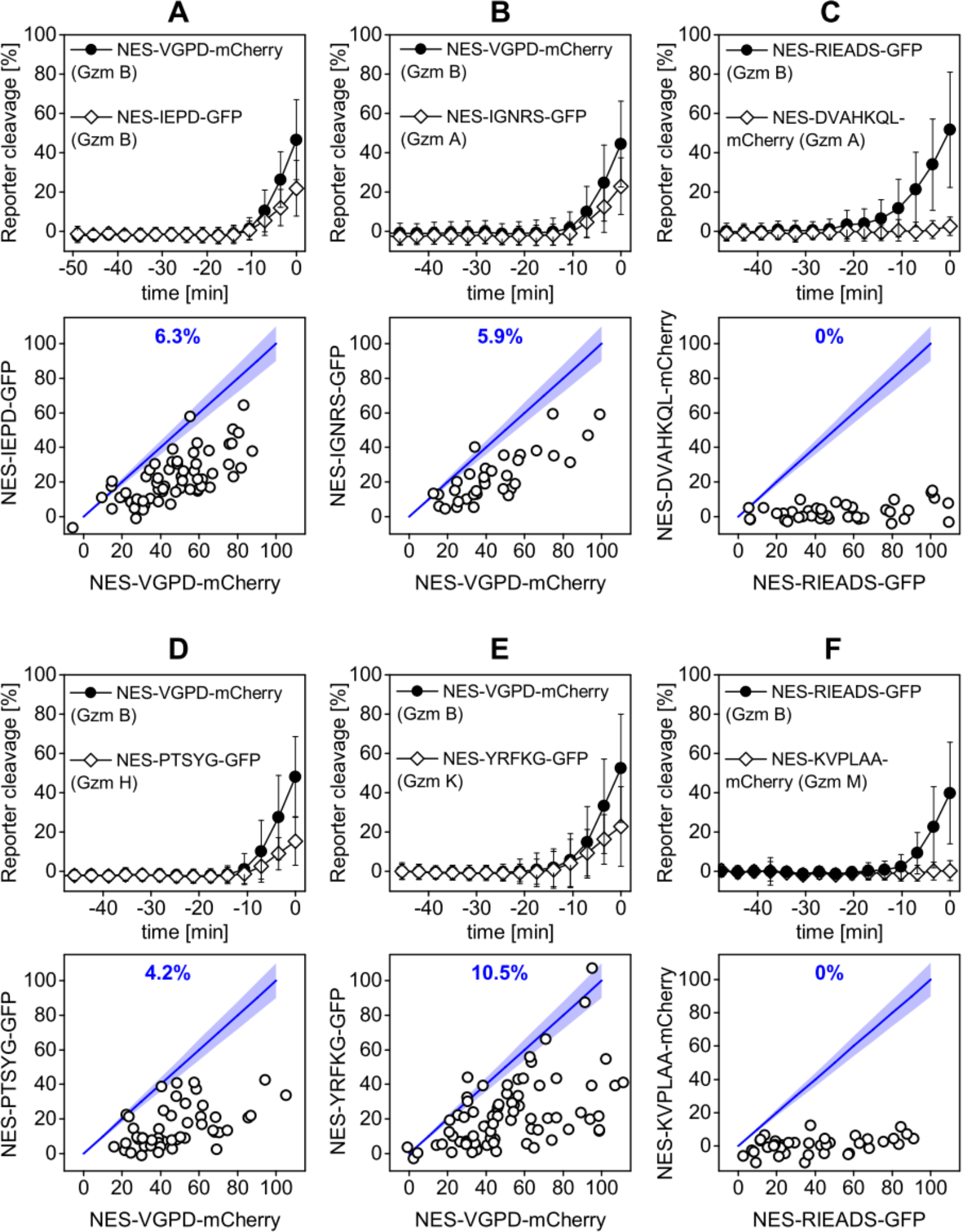
Cleavage of reporters carrying the VGPD and RIEADS cleavage sites for granzyme B is predominant. Quantification of reporter cleavage upon NK92-C1 co-culture (E:T = 3) in HeLa(CD48) cells (A, B, D, E) or HeLa cells transfected with CD48 (C, F), NES-VGPD-mCherry (A, B, D, E) or NES-RIEADS-GFP (C, F) and a second reporter, which was (A) NES-IEPD-GFP for granzyme B, (B) NES-IGNRS-GFP for granzyme A, (C) NES-DVAHKQL-mCherry for granzyme A, (D) NES-PTSYG-GFP for granzyme H, (E) NES-YRFKG-GFP for granzyme K or (F) NES-KVPLAA-mCherry for granzyme M, with 34 to 82 cells per condition. Scatter plots below show reporter cleavage values in single cells at the measurement point before cell death. Percentages (blue) indicate the fraction of single cells showing more or equal (±10%) IEPD-, IGNRS-, DVAHKQL-, PTSYG-, YRFKG- or KVPLAA-reporter cleavage compared to VGPD- and RIEADS-reporter cleavage.

### 4. Discussion

The single-fluorescent protein reporters employed in this study are part of a larger panel of protease reporters, based, for example, on cyclic luciferase (Kanno et al., 2007; Li et al., 2014), fluorophore quenching (Packard et al., 2007), FRET imaging (Choi and Mitchison, 2013; Rehm et al., 2015; Zhu et al., 2016) or subcellular localization of a fluorophore (Beaudouin et al., 2013; Lin et al., 2010). Our reporters provide us with the possibility to precisely investigate NK cell induced cell death since they allow the measurement of two protease activities within the same cell. This feature is useful to correlate the activity of different enzymes within single cells and therefore independently of the large cell-to-cell variability due to the stochasticity of the NK-target cell contact.

We showed that the here developed reporters containing the RIEADS or the VGPD cleavage site are suitable for the measurement of granzyme B activity. We systematically tested putative reporters containing amino acid sequences that were previously shown to be cleaved by granzyme A, B, M, H and K (Mahrus and Craik, 2005). While granzyme B activity was the most prominent, we could detect some activity of granzymes A, H, and K, but no activity of granzyme M. The reporters used here contain a nuclear export signal for the measurement of any enzyme activity in the cytosol. This may explain why the DVAHKQL reporter for granzyme A was not cleaved in target cells upon NK cell addition, since this substrate would naturally reside inside mitochondria (Martinvalet et al., 2008b). However, the contribution of other granzymes to cell death still remains to be further investigated, for example by testing NK cells or CTLs at different maturation stages (Nakazawa et al., 1997) or by taking into account the expression levels over long term experiments (Meiraz et al., 2009).

Our experiments have demonstrated that granzyme B reporters can be applied in combination with caspase-8 or caspase-3 activity reporters. Strong caspase-3 activity correlated with granzyme B activity. These results support earlier studies (Andrade et al., 1998; Pinkoski et al., 2001; Quan et al., 1996; Sutton et al., 2000) showing that granzyme B can directly or indirectly (through BID) induce caspase-3 activation. Our data show that this is a fast process, with caspase-3 activity being detectable within a few minutes after granzyme B activity. In contrast, caspase-8 activity was only detectable in target cells expressing the CD95 receptor, but not in cells where CD95 was knocked-down, despite granzyme B activity. This suggests that there is no major activation of caspase-8 by granzyme B, in contrast to earlier studies (Medema et al., 1997). However, this also demonstrates that NK cells can indeed use two pathways to kill target cells: exocytosis of granzymes and perforin, resulting in detectable granzyme B activity inside the cytosol of target cells, and surface expression of CD95L and TRAIL, resulting in caspase-8 activity. Single-fluorescent protein reporters present a valuable tool to decipher cell death mechanisms induced by NK cells. We believe that this approach opens the door for the characterization of death receptor-versus granzyme-mediated target cell killing, in particular the temporal evolution of these two death mechanisms in the context of serial killing by NK cells.

## 5. Materials and Methods

### 5.1. Constructs

Human CD48 (UniProtKB P09326), human serpin B9/PI-9 (from Gateway cDNA library of the DKFZ) and IZ-sCD95L (Walczak et al., 1999) were subcloned in the pIRES-puro2 vector. Fluorescent reporters were cloned based on constructs described previously (Beaudouin et al., 2013) in pEGFP N1 and C1 vectors (Takara Bio Europe Clontech, Saint-Germain-en-Laye, France). DNA oligonucleotides encoding cleavage sites were cloned with AgeI/NotI or BsrGI/NotI (**Table 2**). The amino acid sequence starting from the N-terminus of the nuclear export sequence (NES) is MNLVDLQKKLEELELDEQQ.

**Table 2:**
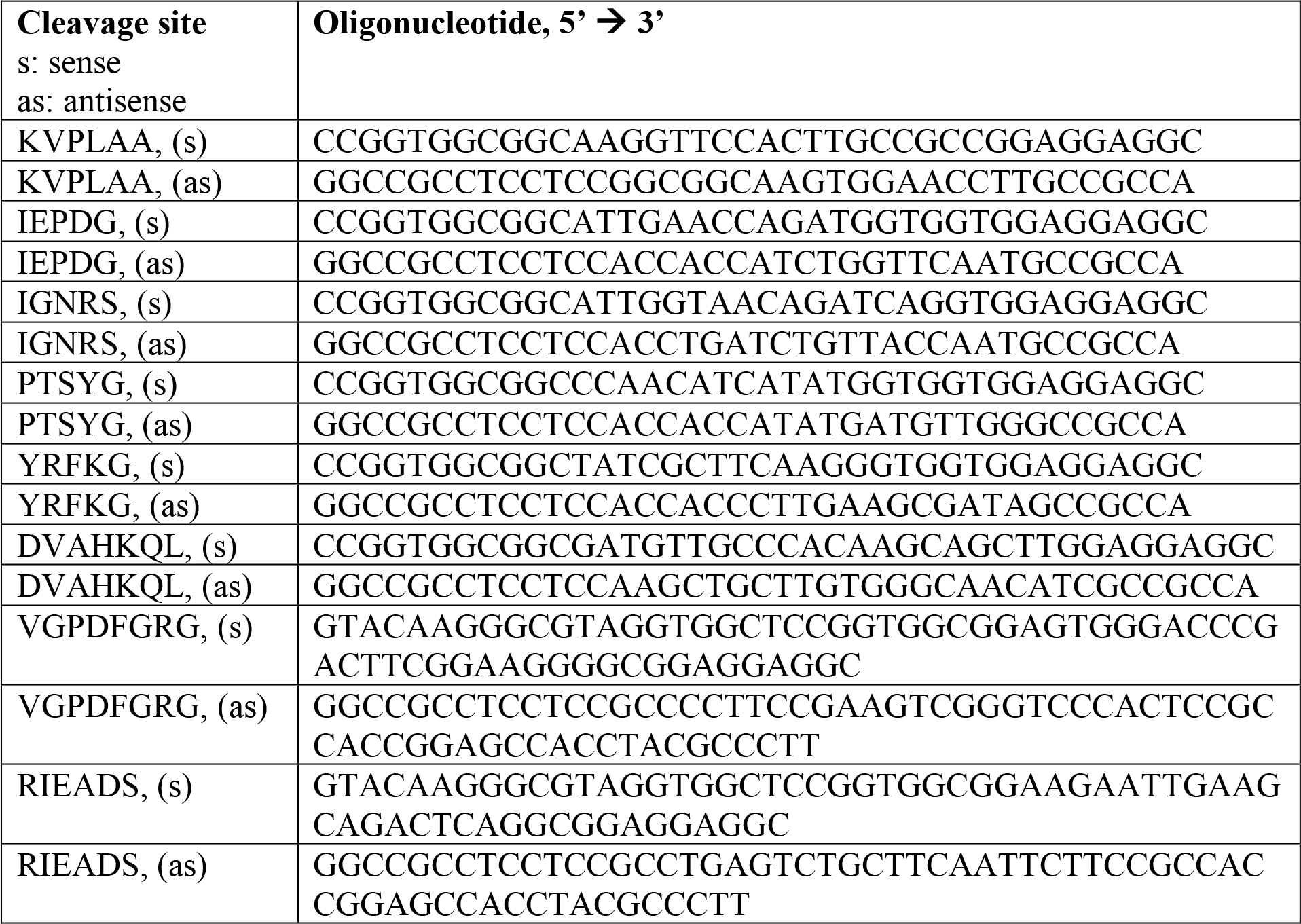
DNA sequence encoding protease cleavage sites used for plasmid cloning.

### 5.2. Cell culture

HeLa cell lines were maintained in Dulbecco’s modified eagle medium (DMEM, Invitrogen, Darmstadt, Germany) containing 10% fetal calf serum (Biochrom AG, Berlin, Germany), penicillin/streptomycin, 100 μg/ml each (Invitrogen). HeLa(CD48) cells over-express CD48 and were maintained in medium supplemented with 0.5 μg/ml puromycin (Sigma-Aldrich). NK92-C1 that stably express IL-2 were maintained in phenol red-free Minimum Essential Medium Eagle Alpha (MEMα, Sigma-Aldrich) without ribonucleosides and deoxyribonucleosides but containing sodium bicarbonate, supplemented with 2 mM L-glutamine, 0.1% 2-mercaptoethanol, 12.5% horse serum, 12.5% fetal bovine serum and penicillin/streptomycin, 100 μg/ml each (Invitrogen).

Soluble CD95 ligand fused to the isoleucine-zipper domain (IZsCD95L, cloned in plRES-puro2) was produced in 293T cells, which were seeded in 6-well plates. 24 h after cell transfection using JetPrime reagent (Polyplus), supernatant was replaced by fresh medium. At the next day, supernatant containing secreted ligand was harvested.

### 5.3. Microscopy

The here presented approach uses fluorescence microscopy, which allows the extraction and correlation of several features, including the time of cell death. Time-lapse microscopy was performed with the TCS SP5 confocal laser scanning microscope from Leica (Leica Microsystems CMS GmbH, Mannheim, Germany) equipped with a 63x/1.4 OIL, HCX PL APO CS objective. We used the live data mode of the Leica software for autofocusing as described in (Beaudouin et al., 2013). The resolution was 512 × 512 pixel and images were acquired in 8-bit or 16-bit. Fluorescence of mGFP and mCherry was acquired in line sequential mode. For mCherry we used the helium-neon laser (561 nm), detection range: 600-660 nm, for mGFP we used the argon laser (488 nm), detection range: 500-560 nm. Cells were imaged in 8-well ibidi chambers (ibidi GmbH, Planegg/Martinsried, Germany). We applied one-, two- or three-fold more effector NK cells compared to target cells (E:T = 1, E:T = 2 or E:T = 3, respectively), hence about 6×10^4^ to 1.8×10^5^ NK cells per well. One field of view in confocal microscopy contained about 2 to 6 transfected cells. All cells expressing both reporters were analyzed. The time-resolution was about 2 to 4 minutes depending on the number of imaged fields and microscopy settings. Equal target cell preparation, equal NK cell preparation (cell number counting, transfection) and parallel measurement of different wells of an 8-well chamber allowed comparison of different conditions in one experiment.

### 5.4. Image analysis

Images were analyzed using ImageJ (Schneider et al., 2012). To quantify the nuclear redistribution of fluorescence intensity over time in single cells, the nuclear intensity was measured. For this, images of each channel were background subtracted. In most cases, two reporters were measured within one cell: to analyze them, one channel was assigned green, the other red, both were transformed into RGB, and they were then superimposed. The mean intensity within the nucleus was quantified by choosing a representative region within the nucleus at each time point until cell rounding due to cell death was observed. We chose to normalize the nuclear intensity to the cytosolic intensity at each time point for soluble-cytosolic reporters to calculate the percentage of reporter cleavage.

### 5.5. Statistics

Data were visualized and analyzed using OriginLab software. Data groups were considered significantly different as indicated in the text, but considered not significantly different when P values were greater than 0.05. Boxplots additionally show raw data points. Box and lines indicate median, 25% quartile and 75% quartile. To plot the mean kinetics of reporter cleavage of single cell data, we set the time point that directly preceded cell death to time = 0 and calculated the mean and standard deviation of single cell responses. The time values can therefore have a negative sign on the plots.

## Acknowledgements

We kindly thank the DKFZ for providing the Gateway cDNA sequence of human serpin B9/PI-9. This work was supported by the Initiative and Networking Fund of the Helmholtz Association within the Helmholtz Alliance on Systems Biology/ SBCancer. CW is supported by funding of the Leibniz Association (SAW-2013-IfADo-2) and the Deutsche Forschungsgemeinschaft (DFG, WA-1552/5-1).

## Author contributions

CL designed the study, performed experiments, analyzed the data and wrote the paper. PS performed experiments and analyzed data. MC isolated and purified primary human NK cells. RE, JB and CW supervised the work and helped writing the paper.

